# Age-Related Alterations in Multispectral Somatosensory Gating: Evidence for Partial Compensation in Attentional Performance

**DOI:** 10.64898/2026.06.05.730439

**Authors:** Mahak Virlley, Yin Xi, Natalie M. Bell, Tyrell Pruitt, Lin Guo, Sloan White, Fang F. Yu, Laura H. Lacritz, Heidi Rossetti, C. Munro Cullum, Amil M. Shah, Elizabeth M. Davenport, Joseph A. Maldjian, Amy L. Proskovec

## Abstract

Healthy cognitive aging involves selective changes, with relative preservation of some domains and decline in others, particularly attention, inhibitory control, and executive function. Somatosensory gating (SG) refers to the brain’s ability to suppress neural responses to redundant tactile input, conserving resources for relevant stimuli and reflects pre-attentive and inhibitory mechanisms. Prior region-of-interest studies have shown age-related reductions in gamma SG within contralateral primary somatosensory cortex (S1), and modulation of theta, alpha, and beta SG by attention in young adults. However, whole-brain, multispectral age effects remain unclear. In this study, 63 middle-to-older aged adults (38 females; mean age = 59.9 ± 8.6 years) underwent magnetoencephalography during a paired-pulse somatosensory paradigm. SG was quantified as attenuation of the neural response to the second stimulus relative to the first. Time-frequency analyses identified theta (4-7 Hz), alpha (8-13 Hz), beta (15-25 Hz), and gamma (30-90 Hz) oscillatory responses, and band-specific voxel-wise whole-brain gating maps assessed age-related effects. Attention/executive function was also measured. Results showed significant age-related increases in gamma SG in the contralateral supplementary motor area. Mediation analyses suggested this increase partially offsets age-related declines in attention/executive function, consistent with a partial compensatory mechanism. Additionally, theta SG in contralateral S1 increased with age. These findings demonstrate frequency- and region-specific age-related alterations in SG, suggesting that older adults may recruit enhanced inhibitory mechanisms, particularly in higher-order sensorimotor regions, to support cognitive function.

## Introduction

Healthy cognitive aging is susceptible to variable changes in cognition, such as declines in attention span and capacity to inhibit distractions [1, 2]. Of interest, somatosensory gating (SG) is a neural inhibitory mechanism in which the brain filters out redundant tactile stimuli from the surrounding environment in a time-dependent manner to conserve neural resources for processing behaviorally-pertinent information [3]. Typically, when two coupled and identical tactile stimuli are issued rapidly (~500 ms separation), the cortical response to the second stimulus is attenuated, reflecting efficient inhibitory processing [3–7]. SG has been shown to be disrupted in neurodegenerative, neuropsychiatric, and sensorimotor conditions, underscoring its importance for both behavior and cognition [8–17]. In healthy populations, SG is most consistently observed in the gamma frequency band (30+ Hz) within the primary somatosensory cortex (S1), and evidence suggests that gamma SG diminishes with advancing age, indicating reduced inhibition of redundant stimuli [6, 18]. However, prior studies have largely relied on region-of-interest (ROI) approaches, predominantly focusing on S1, which may obscure broader network-level changes. Given that SG likely reflects distributed inhibitory processes across multiple cortical regions [14], a whole-brain, voxel-wise approach is warranted to more comprehensively characterize age-related changes in SG.

Beyond gamma activity, SG has also been observed in lower frequency oscillations, including theta (4-7 Hz), alpha (8-14 Hz), and beta (15-29 Hz), which are implicated in somatosensory and cognitive processing [4, 19–22]. Oscillatory dynamics in these bands are thought to capture distinct neural mechanisms not reflected in time-domain responses, particularly those related to large-scale network coordination and cognitive control [4, 19, 20]. For instance, theta, alpha, and beta SG within S1 are modulated by attentional demands [4]. Specifically, enhanced suppression of the second stimulus in the theta band occurs when attention is directed toward somatosensory input, reflecting amplified theta SG under increased attentional engagement [4]. Despite this, age-related SG studies have primarily focused on gamma activity [5, 6, 14, 17, 18], leaving a critical gap in understanding how lower-frequency oscillatory gating may change with age.

Importantly, SG is closely linked to attention/executive function processes, as it reflects the brain’s ability to suppress irrelevant sensory input while prioritizing salient information. Attentional subdomains, including sustained attention and inhibitory control, degrade with age [1, 23–25], suggesting that age-related alterations in SG may contribute to these cognitive changes. For example, regions such as the supplementary motor area (SMA) are implicated in “gating” of basic sensory perception [26], yet their role in SG remains underexplored. SMA is thought to process higher-order cognitive components along with sensorimotor control, such as motor execution [26–28]. Investigating SG across both primary and higher-order regions may therefore provide insight into how distributed neural systems support attention/executive function in aging.

To address these gaps, the present study applied a whole-brain, voxel-wise approach to examine SG across multiple frequency bands in a middle-to-older aged population. We generated band-specific voxel-wise gating difference maps (Stim1–Stim2) to characterize age-related changes in SG and evaluated their relationship with attention/executive function. We hypothesized that age-related alterations in SG would emerge across both theta and gamma frequency bands within sensorimotor and higher-order cortical regions. Specifically, we predicted that theta SG in contralateral S1 would increase with age. We further hypothesized that gamma SG in higher-order sensory regions would be enhanced with age. Importantly, we expected that both theta SG in S1 and gamma SG in higher-order regions would be associated with attention/executive function, reflecting compensatory mechanisms that support cognitive functioning in healthy aging.

## Methods

### Participants, study, and ethics

The study utilized data collected as part of the Dallas Hearts and Minds Study (DHMS) [29], which was reviewed and approved by the Institutional Review Board at the University of Texas Southwestern Medical Center. The DHMS is a large longitudinal population-based study enriched for racial minorities with a focus on cardiovascular and neurocognitive health. Written informed consent was obtained from each participant following a detailed description of the study protocol. Pertinent to this study, data collected included demographics, MRI, medical history forms, a neurocognitive assessment battery, and, in a subset of participants, MEG. MEG was collected in a subset of participants due to scheduling constraints on the machine (i.e., mornings were devoted to clinical epilepsy patient scans), and exclusion of participants with ferromagnetic implants which are contraindicated (i.e., cochlear implants, magnetic shunts) or caused significant artifact in the MEG signal (e.g., irremovable dental bridge). As the present study was focused on healthy cognitive aging, additional exclusionary criteria included neurological or current neuropsychiatric disorders, concussion history within the past year, current high intensity alcohol use [30], or any other medical illness affecting the central nervous system. Cognitive status (e.g., non-impaired, cognitively impaired) of each participant was determined by consensus amongst two board-certified neuropsychologists who reviewed demographic, medical history, and neurocognitive data. Only participants designated as non-impaired were included in the present study. Additionally, as presence of subjective cognitive complaints may be indicative of preclinical Alzheimer’s disease and related dementias, we further excluded participants who endorsed subjective cognitive complaints from the analysis [31, 32]. Specifically, participants who responded “yes” to items inquiring “Do you have any problems with your memory?” and/or “Do you have any difficulties solving problems?” were excluded.

### Neurocognitive measure of attention/executive function

The DHMS cognitive battery was administered by trained research personnel and included standardized measures of global cognition and domain-specific performance. Please see the published DHMS protocol for a complete description of the neurocognitive assessment [29]. Relevant to this study, Trail Making Test Parts A and B (TMT-A/B), which assesses attention and processing speed (TMT-A) and executive function, inhibitory control, and divided attention (TMT-B), were administered [33–35]. Participants were asked to connect a sequence of circled items as quickly as possible without lifting their pencil. For part A, on a sheet containing numbers (1-25), participants were instructed to connect the circles in ascending order. For part B, on a sheet containing numbers (1-13) and letters (A-L), participants were instructed to alternate between the two sequences in ascending order (1 → A → 2 → B → 3 → C, etc.). Completion time (in seconds) served as the primary outcome, with longer times indicating slower processing or reduced executive control. Errors were immediately corrected by the examiner and contribute to overall completion time. Due to previous studies linking somatosensory processing and SG with TMT-A/B [17, 36], TMT-A/B completion time was initially selected as the primary index of attention/executive function. However, only TMT-B showed a significant association with SG; therefore, subsequent analyses were restricted to TMT-B. Due to past literature suggesting race, education, and sex have an impact on neurocognitive test performance [37, 38], raw neurocognitive scores were entered into linear regression models to regress out the effects of race, education, and sex using R [39]. Residualized TMT-B time was utilized in final analyses and will be referred to as “TMT-B time” henceforth.

### Experimental paradigm

During MEG, participants were placed in the 60° seated position in a nonmagnetic chair with an accompanying tray and their head touching the top of the MEG helmet sensor array. Additionally, foam padding was placed around the head to minimize head motion, and they were instructed to close their eyes and relax their hand and arm for the task. Unilateral electrical stimulation was applied to the right median nerve using external cutaneous stimulators with a Digitimer DS7A constant-current stimulator system. 100 paired-pulse trials were collected with an interstimulus interval of 500 ms and inter-pair interval of 3950 ± 150 ms. Thus, a total of 200 stimuli were delivered during the task resulting in a total run-time of ~7 minutes. Each pulse generated a 0.2 ms constant current square wave that was set to 10% above the motor threshold required to elicit a subtle thumb twitch. The stimulation level for each participant was recorded and included as a covariate in regression analyses; it was only reported when significant covariance was observed.

### MEG data acquisition

Recordings were conducted using a 306-sensor MEGIN TRIUX Neo MEG system, containing 204 planar gradiometers and 102 magnetometers, at a sampling rate of 1 kHz and acquisition bandwidth of 0.1-330 Hz in a 3-layer magnetically-shielded room. Participants were monitored for safety using a real-time audio-video system. Preceding MEG, a 3D digitizer was used to create a map of the subject’s head including the location of fiducials and head position indicator coils. Further, during the MEG recording, an electric current with a unique frequency (e.g., 320 Hz) was fed to each coil, inducing a magnetic field allowing each coil to be continuously localized with reference to the sensors, and head motion to be corrected for offline. Following the recording, MEG data was noise reduced and corrected for head motion using the signal space separation method with a temporal extension [40–42].

### Structural MRI acquisition, processing, and MEG coregistration

Following the MEG scan, fiducial markers that appear hyperintense on T1-weighted MRI were placed at the nasion and left and right preauricular areas marked during the MEG digitization step. These markers remained in place until the completion of the MRI to aid in co-registration. Individual structural MRI data were obtained using a 3T Siemens Prisma scanner with a 64-channel head coil. A whole-brain 3D T1-weighted Magnetization-Prepared Rapid Gradient Echo (MP-RAGE) sequence with 1-mm^3^ isotropic resolution was acquired using the following parameters: repetition time, 1800 ms; echo time, 2.26 ms; field of view, 256 mm; matrix, 256 x 256; slice thickness, 1mm; acquisition plane, sagittal; flip angle, 8 degrees. Structural MRI data were processed using the FreeSurfer (version 7.1.1) recon-all pipeline. The segmented MRI data were then imported into Brainstorm (Neuroimage, USC; version 3.4) [43]. Since head position indicator coil locations were known in head coordinates, MEG data were transformed into a common coordinate system. Using this common coordinate system, each participant’s MEG data were co-registered to their structural T1-weighted brain MRI prior to source space analyses. Following source analysis, each participant’s functional images were transformed into standardized MNI space [44] using the transform applied to the structural MRI data. Structural and diffusion MRI analyses were also conducted to examine potential associations with SG; detailed methods (including diffusion MRI acquisition) are provided in the Supplementary Information.

### MEG preprocessing

All MEG data underwent standard preprocessing detailed in the DHMS MEG protocol manuscript [29]. A high-pass filter of 1 Hz, low-pass filter of 200 Hz, and notch filter at 60 Hz and harmonics (i.e., 120 Hz, 180 Hz) were applied to the MEG data. Artifacts (e.g., cardiac, ocular) were removed using independent component analysis. MEG data were visually inspected for additional artifact contamination (e.g., muscle), and those segments with remaining artifact were rejected. Cleaned, continuous data was split into epochs assigned as 4300 ms in length with 0 ms denoting delivery of the first stimulus and 500 ms denoting the delivery of the second stimulus. Baseline was defined as −700 to −300 ms. An average of 87 (SD = 8) trials per participant were used for analysis. The number of usable trials did not vary by age, (*p* > 0.05, linear regression). Only data from planar gradiometers were used during subsequent analyses.

### MEG sensor-level analysis

Artifact-free epochs were transformed into the time frequency domain using Morlet wavelet analysis (range: 2-152 Hz, step: 1 Hz), averaged across trials to generate time-frequency plots of mean spectral density per sensor, and subsequently baseline-normalized. The time-frequency windows used for imaging were determined by statistical analysis of the sensor-level spectrograms across all participants using a two-stage procedure. First, parametric tests against a null hypothesis of a zero mean were conducted and the output spectrograms of t-values were thresholded at *p* < 0.05. Next, multiple comparisons were controlled for using the Benjamini-Hochberg false discovery rate (FDR) procedure. Only time frequency windows that contained significant oscillatory responses across participants were subjected to beamforming (i.e., imaging) analysis.

### MEG source-level analysis

Cortical networks were imaged via a linearly constrained minimum variance vector beamformer, which calculates source power for the entire brain volume by employing spatial filters in the time-frequency domain [45–47], and normalized for each participant using a separately averaged pre-stimulus noise period (i.e., baseline) of equal duration and bandwidth [45]. These images are commonly referred to as pseudo *t*-maps, with units (pseudo-*t*) that reflect noise-normalized (i.e., relative) power per voxel at a 4.0 × 4.0 × 4.0 mm resolution. The maps were transformed into standardized MNI space and spatially resampled to 1 mm^3^ voxels. Two participant-level whole-brain maps were generated (i.e., Stimulation 1 [Stim1] and Stimulation 2 [Stim2] in the paradigm) for each oscillatory response identified in the sensor-level analysis. Whole-brain gating difference (GD) maps were created for each oscillatory response per participant by subtracting the Stim2 map from the Stim1 map (Stim1-Stim2). These GD maps indexed each participant’s ability to gate the second stimulus in the pair. All MEG pre-processing and imaging took place in Brainstorm (version 3.4) [43].

### Statistical analyses

The CAT12 toolbox in Statistical Parametric Mapping (SPM12) was employed for whole-brain regression analyses [48]. To identify regions in which SG varied significantly with age, the whole-brain GD maps were entered into a linear regression with participant age. The resulting statistical parametric maps were displayed as a function of alpha level = 0.005 and adjusted for multiple comparisons using a cluster threshold (*k* = 500 contiguous voxels) based on Gaussian random field theory [49–51]. As described in *Results*, significant age effects were only present in theta and gamma frequency bands and, therefore, subsequent analyses focused exclusively on these bands.

To further investigate age-related effects, voxel time-series data (“virtual sensors”) were extracted from each participant’s MEG data using the peak voxel coordinates from each region displaying significant GD age effects. Virtual sensors were generated by applying the sensor weighting matrix obtained from the forward model to the preprocessed signal vector, which produced a time series corresponding to each data-driven region-of-interest (ROI). These data were decomposed into time-frequency space following the time-frequency extents used in the beamformer analysis for each oscillatory response. The primary relationships of interest were those between age and the gating ratio (GR) and spontaneous power for theta and gamma in their respective ROIs. From the relative (i.e., baseline-normalized) time series we extracted the peak response amplitudes following Stim1 and Stim2 per participant and used these values to compute the GR (Stim2/Stim1) per participant. Of note, GR was of interest as it accounts for overall power, whereas the GD maps used in the whole-brain voxel-wise analyses did not. To avoid negative GR values and dividing by values close to zero, the least common number was added per frequency band. Specifically, the number 6 was added to offset negative theta response amplitudes, and the number 8 was added to offset negative gamma response amplitudes. Of note, values were added to gamma and theta because both frequency bands demonstrate event-related synchronizations. GR was calculated using these offset response amplitudes per participant. From the absolute (i.e., not baseline-normalized) time series we extracted spontaneous amplitude per participant by averaging across the baseline time window. For each ROI two simple regressions were computed in which GR and spontaneous amplitude at the relevant frequency band were independently regressed on age. Secondary analyses independently regressed Stim1 and Stim2 response amplitude on age, per ROI. To evaluate neurobehavioral relationships, analyses were performed integrating neurocognitive and MEG-derived metrics. As mentioned in the Introduction, we hypothesized that inter-individual variability in functional inhibition (i.e., SG) may hold relevance for age-related declines in attention/executive function. Thus, additional linear regression models assessed: (1) the effect of age on TMT-B time, (2) the effect of MEG metrics on TMT-B time, and (3) the association between age and TMT-B time controlling for MEG metrics. To determine whether altered somatosensory responses mediated age-related cognitive deficits, mediation analyses were performed testing the pathway: Age → SG → TMT-B time [52, 53]. Indirect (Average Causal Mediated Effect; ACME) and direct (Average Direct Effect; ADE) effects were estimated using nonparametric bootstrapping (10,000 iterations) with bias-corrected 95% confidence intervals implemented via the R *mediation* package [54]. A significant indirect effect (i.e., ACME) was interpreted as evidence that somatosensory responses mediated the relationship between age and cognitive performance. All neurobehavioral statistical analyses were performed in R using RStudio [39].

## Results

In total, 319 DHMS participants completed the MEG SG paradigm. Of those, 116 were cognitively healthy (CH) (i.e., absence of objective and subjective cognitive deficits). From this CH sample, 42 participants were disqualified from this study for medical morbidities (e.g., neurological disorder). One participant was scanned using the left hand for SG due to an injury; this participant’s data was not used. Four participants were omitted due to incomplete demographic data. Lastly, three participants’ MEG data was unusable due to the presence of excessive artifact, and three others qualified as MEG metric outliers (i.e., greater than three standard deviations beyond the mean). After removal of these individuals, the data from 63 participants were used in final analyses (Table 1). The demographics (sex, race, and education) and MoCA scores were not related to aging (p > 0.05) and thus, were not evaluated further in the context of the healthy aging analyses. MoCA scores fall under normative standards for an ethnically diverse population [55].

**Table 1.** Demographics for a subset of DHMS participants that qualified for the present study design.

| Number of participants | Age | MoCA | Sex | Race | Education |
| --- | --- | --- | --- | --- | --- |
| 63 | Average ± Standard Deviation:<br>59.9 ± 8.6 years old<br>Median: 61 years old | Average ± Standard Deviation:<br>25 ± 3<br>Median: 25 | 38 female<br>25 male | 23 Black or African American<br>30 White or Caucasian<br>2 American Indian<br>2 Asian<br>3 Other<br>2 Multiple Races<br>1 Unknown | Average ± Standard Deviation: 15 ± 2 years<br>Median: 16 years |

### Neural oscillatory responses to the SG paradigm

Following each stimulation, robust increases in theta (4-7 Hz, 0-250 ms post stimulus; *p* < 0.05 corrected) and gamma (30-90 Hz, 10-60 ms post stimulus; *p* < 0.05, corrected) activity were observed (Figure 1A). Further, decreases (i.e., event-related desynchronizations) in beta (15-25 Hz, 125-350 ms post stimulus; *p* < 0.05, corrected) and alpha (8-13 Hz, 225-400 ms post stimulus; *p* < 0.05, corrected) activity were found (Figure 1A). These responses were largely captured in left parietal and posterior frontal sensors near the sensorimotor cortices. Thus, we applied a beamformer to the following windows: 4-7 Hz from 0-250 ms (Stim1) and 500-750 ms (Stim2), 30-90 Hz from 10-60 ms (Stim1) and 510-560 ms (Stim2), 15-25 Hz from 125-350 ms (Stim1) and 625-850 ms (Stim2), and 8-13 Hz from 225-400 ms (Stim1) and 725-900 ms (Stim2). All oscillatory responses localized to the hand-knob region of contralateral S1. As exemplified in Figure 1B, the resulting whole-brain maps were used to compute GD (Stim1 – Stim2) maps for each frequency-specific response, per individual.

**Figure 1.**
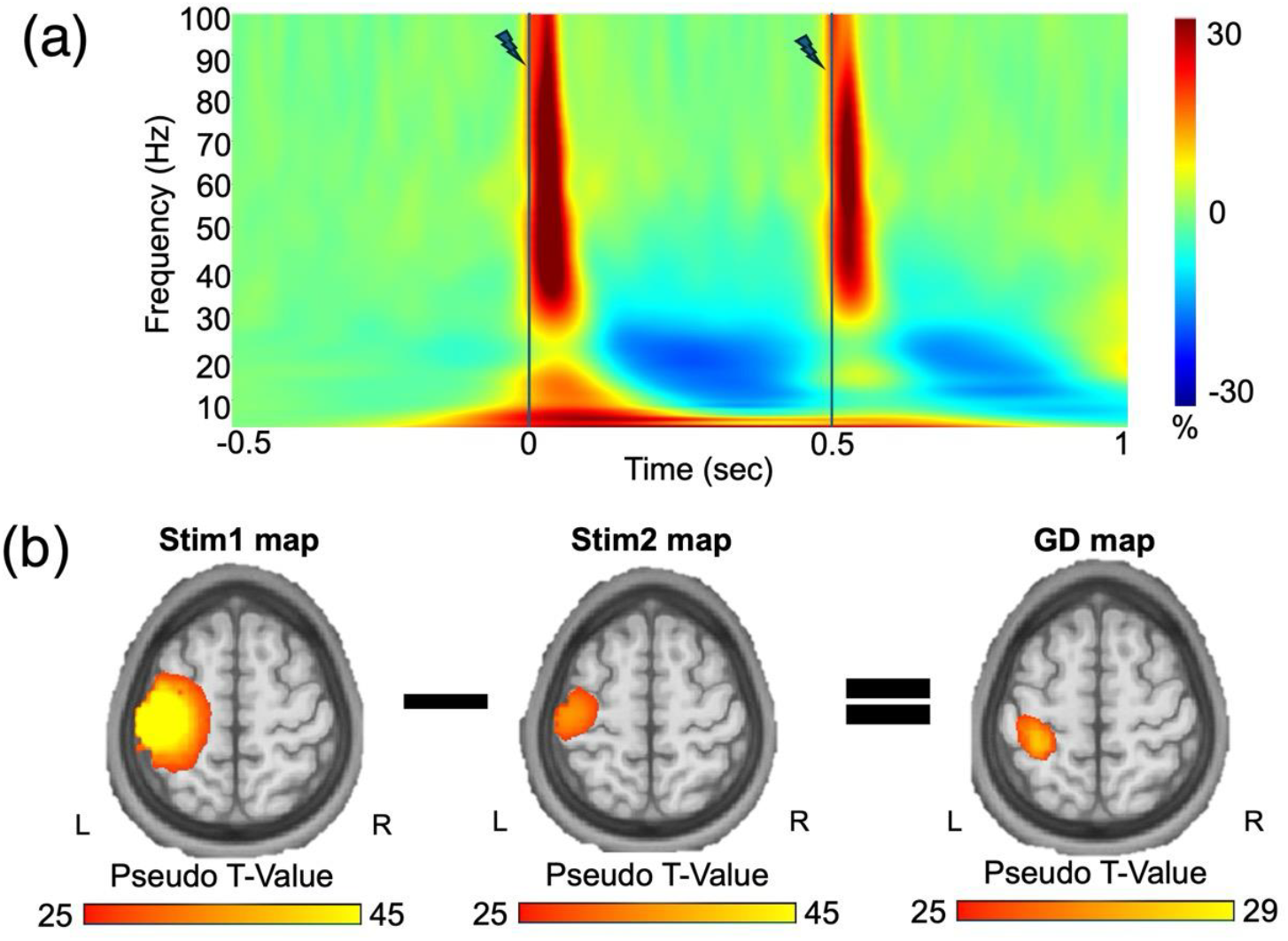
Whole-brain Somatosensory Gating Analysis Pipeline. (a) Grand-averaged time-frequency spectrogram from a peak sensor near the contralateral sensorimotor cortex. The x-axis denotes time in seconds, beginning at −0.5 seconds and ending at 1 second. Stimulation was delivered at 0 seconds and 0.5 seconds, denoted by the line and bolt symbol. The y-axis denotes frequency (Hz), and the magnitude is shown as a percentage change unit relative to the baseline period, with a color bar to the far right of the spectrogram. As can be discerned, strong theta, gamma, beta, and alpha responses followed each stimulation. These sensor-level data were utilized to define the time-frequency windows subjected to beamforming. (b) Theta (4-7 Hz) beamformer maps (pseudo-t values) for Stim1 and Stim2, as well as the gating difference (GD) map are shown for a representative participant. The GD map was calculated as Stim1 – Stim2 across each voxel within the 3D whole-brain map. GD maps were computed in a similar fashion for gamma, beta, and alpha per participant (not shown).

### Theta SG and Spontaneous Theta in Contralateral S1 as a function of age

To identify brain regions exhibiting changes in SG as a function of age, we performed whole-brain regressions using the GD maps as the dependent variable and participant age as the independent variable, per oscillatory response. As hypothesized, a significant age effect was observed for theta SG in the hand-knob region of contralateral S1 (*p* < 0.005, corrected; Figure 2A). To ensure that the effect was not due to age-related variance in overall power, relative time series were extracted from the peak voxel within this region (Figure 2B) and used to calculate the GR by dividing Stim2 response amplitude by Stim1 response amplitude (Stim2/Stim1) per participant. We found that contralateral S1 theta GR significantly decreased with age (*β* = −0.355, *t*(61) = −2.962, *p* = 0. 004), indicative of exaggerated SG with advancing age (i.e., greater functional inhibition with age; Figure 2C). No significant age effects were observed for relative theta Stim1 or Stim2 response amplitude within contralateral S1 (*p*’s > 0.05).

**Figure 2.**
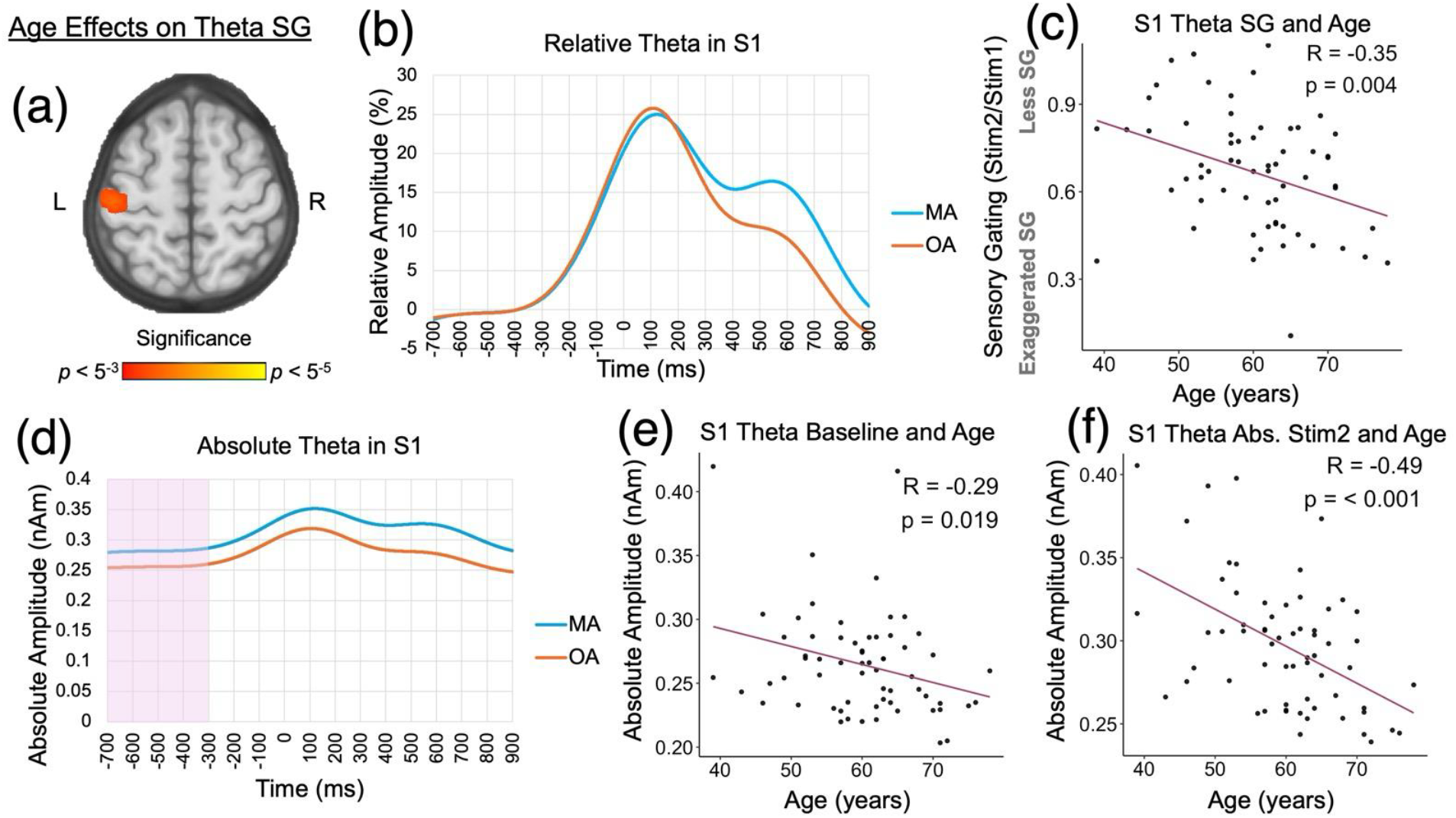
Age Effects on S1 Theta SG. (a) A cluster with peak voxel located in the hand region of contralateral S1 was observed after regressing theta (4-7 Hz) SG maps on age, indicating a significant age effect on theta gating within the region (*p* < 0.005, corrected). (b) To visualize age effects on SG, we split participants into two groups consisting of middle-aged (MA) adults (blue; age range: 39-54 years old, N = 17) and older-aged (OA) adults (orange; age range: 65-78 years old, N = 17); individuals falling within ±0.5 SDs of the entire sample’s average age were excluded, those below 0.5 SDs of the group’s mean comprise the MA adults, and those above 0.5 SDs of the group’s mean comprise the OA adults. Age-related gating differences are clearly observed in the theta relative amplitude time series extracted from the peak voxel in contralateral S1. (c) Age was negatively associated with theta SG ratio, reflecting a lower gating ratio (i.e., exaggerated gating) in OA adults. (d) Theta absolute time series displayed in a similar fashion as panel b revealed depressed spontaneous activity (baseline amplitude) in contralateral S1 for OA adults. The baseline period (−700 ms to −300 ms) is highlighted in pink. (e) Age was negatively correlated with absolute amplitude during the baseline period, such that, as age increased, spontaneous theta decreased. (f) Age was negatively correlated with absolute theta amplitude following Stim2, such that, as age increased, absolute theta Stim2 response amplitude in contralateral S1 decreased. This effect was also found beyond the effects of baseline (see Supplemental Figure 1 for residual plot). S1 = primary somatosensory cortex. SG = somatosensory gating.

To investigate the relationship between spontaneous theta within contralateral S1 and age, the absolute time series was extracted from the peak voxel (Figure 2D) and used to compute the average absolute amplitude during baseline per participant. Spontaneous theta within contralateral S1 decreased significantly with advancing age (*β* = −0.295, *t*(61) = −2.407, *p* = 0.019; Figure 2E). We additionally probed absolute theta Stim1 and Stim2 response amplitude and absolute GR. Absolute theta Stim1 response amplitude did not correlate with age, irrespective of whether baseline was accounted for (*p*’s > 0.05). Absolute theta Stim2 response amplitude negatively correlated with age, both when baseline was not accounted for (*β* = −0.489, *t*(61) = −4.373, *p* < 0.001; Figure 2F), as well as beyond the effects of baseline (*β* = −0.292, *t*(60) = −3.629, *p* < 0.001; Supplemental Figure 1A). Finally, absolute theta GR within contralateral S1 significantly decreased with advancing age (*β* = −0.435, *t*(61) = −3.775, *p* < 0.001).

### Gamma SG and Stim1 amplitude in Contralateral SMA as a function of age

The whole-brain voxel-wise analysis for gamma GD revealed a significant age effect for gamma SG in the contralateral SMA (*p* < 0.005, corrected; Figure 3A). Utilizing the same analytical pipeline as that described for theta in section 3.3, we found that contralateral SMA gamma GR significantly decreased with age, indicating exaggerated SG with advancing age in this region (*β* = −0.387, *t*(61) = −3.28, *p* = 0.002; Figure 3C). In addition, relative gamma Stim1 response amplitude in contralateral SMA was positively correlated with age (*β* = 0.357, *t*(61) = 2.989, *p* = 0.004; Figure 3D). However, an age effect was not found for relative gamma Stim2 response amplitude (*p* > 0.05).

**Figure 3.**
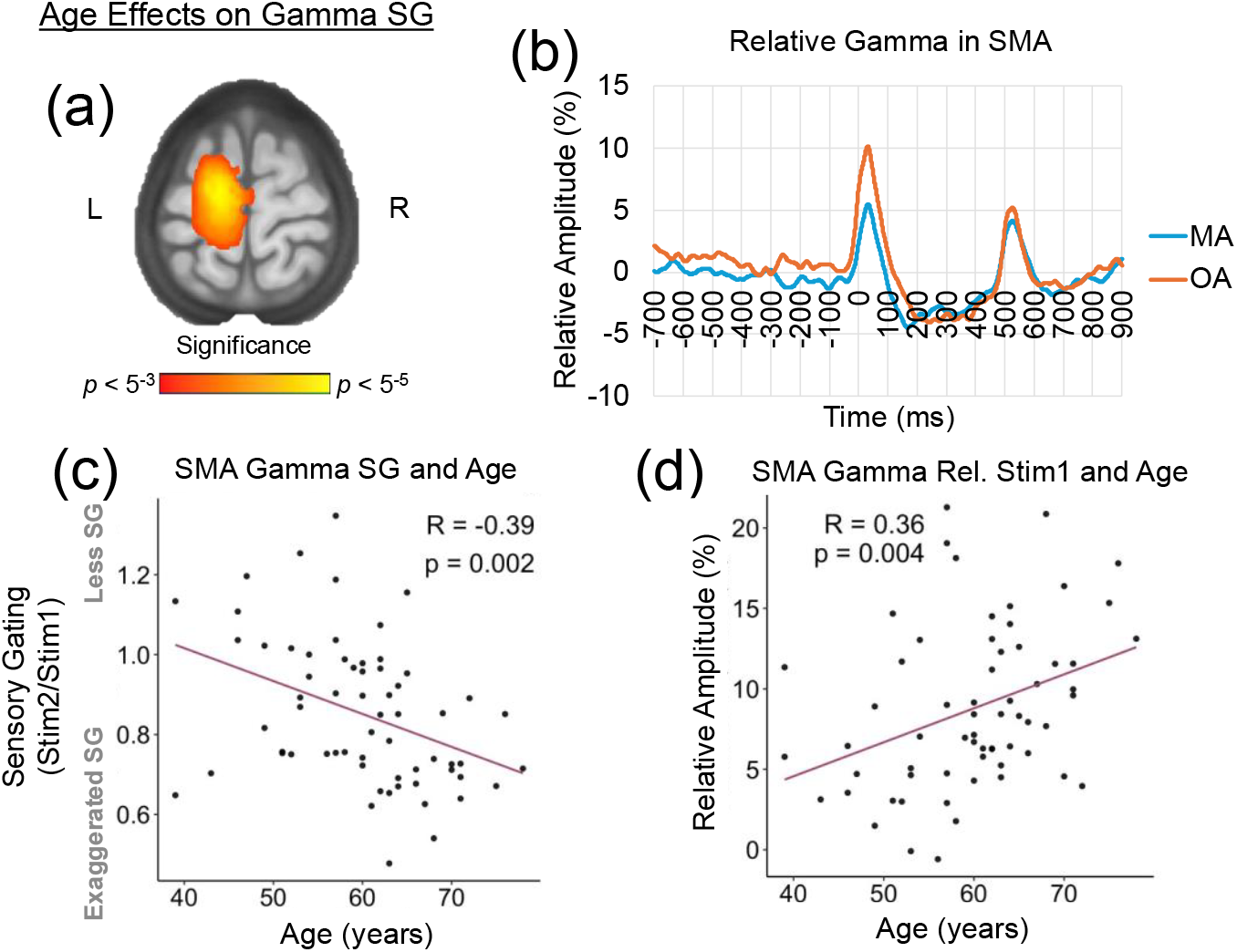
Age Effects on SMA Gamma SG. (a) A cluster with peak voxel located in contralateral SMA was observed after regressing gamma (30 - 90 Hz) SG maps on age, indicating a significant age effect on gamma gating within the region (*p* < 0.005, corrected). (b) When separated into MA and OA adults following the 0.5 SD method (see Figure 2 caption for details), age-related gating differences are clearly observed in the gamma relative amplitude time series extracted from the peak voxel in contralateral SMA. (c) Age was negatively correlated with gamma SG ratio, reflecting a lower gating ratio (i.e., exaggerated gating) in OA adults. (d) Age was positively correlated with relative gamma Stim1 response amplitude such that, as age increased, gamma Stim1 response amplitude in contralateral SMA also increased. SMA = supplementary motor area. SG = somatosensory gating. MA = middle-aged. OA = older-aged.

Regarding the absolute gamma metrics, the results indicated that contralateral SMA absolute GR significantly decreased with advancing age (*β* = −0.429, *t*(61) = −3.704, *p* < 0.001). No significant age effects were observed for absolute Stim1 response amplitude, absolute Stim2 response amplitude, or baseline absolute amplitude for gamma in contralateral SMA (*p*’s > 0.05).

Although the whole-brain voxel-wise analysis demonstrated additional age-related changes in gamma SG in the right medial temporal gyrus and right inferior frontal gyrus (*p*’s < 0.005), these regions showed effects limited to the GD measure and did not demonstrate corresponding age-related changes in gamma GR (*p*’s > 0.05). As these age effects did not survive after accounting for differences in overall response power and given that the GR is the most consistently accepted metric of sensory gating, additional probing of these regions was not pursued.

No significant age effects were found for beta or alpha SG through the whole-brain voxel-wise GD analysis.

### SMA Gamma SG Exaggeration Partially and Inconsistently Mediates Attention/Executive Function Performance with Age

To further investigate potential implications of exaggerated SG within the context of cognitive aging, we assessed whether SG contributed to age-related cognitive slowing using TMT-B time (demographic-adjusted residuals) as a measure of attention/executive function. Longer time indicated worse cognition, specifically attention/executive function. Replicating previous studies [23, 24], a simple linear regression revealed that age was a significant predictor of TMT-B time, such that older age was associated with slower TMT-B performance (*β* = 0.389, *t*(61) = 3.298, *p* = 0.002; Supplemental Figure 2A). A multiple linear regression revealed that age effects on TMT-B time were greater after inclusion of contralateral SMA gamma SG in the model (β = 0.512, t(60) = 4.184, p < 0.001; Supplemental Figure 2B). This model also revealed that exaggerated SMA gamma SG was a significant predictor of faster TMT-B time (i.e., better attention/executive function; *β* = 0.317, *t*(60) = 2.592, *p* = 0.012; Supplementary Figure 2C). Causal mediation analysis (10,000 bootstrap samples) supported a significant indirect effect of age on TMT-B time through SMA gamma SG (ACME = −0.123, 95% CI [−0.304, −0.018], *p* = 0.009; Figure 4). The indirect effect was negative, indicating that age-related exaggeration of SG partially offset age-related attention/executive function decline. The negative indirect effect indicates that age-related exaggeration of SG partially compensated for cognitive decline; however, controlling for this compensatory response unmasked a stronger direct effect of age on attention, consistent with statistical suppression. The direct effect of age (ADE = 0.512, 95% CI [0.278, 0.802], *p* < 0.001) and the total effect (0.389, 95% CI [0.0161, 0638], *p* < 0.001) were both significant (Figure 4), with the former indicating that compensatory changes in gating were insufficient to overcome the broader detrimental effects of aging on cognition. The direct and indirect effects have opposite signs, and the inclusion of the mediator variable increased the variance between age and TMT-B time, indicating inconsistent mediation [56]. About 32% of the total effect of age on attention/executive function was mediated by gamma SG (p = 0.010), which reflects partial mediation (Figure 4). Taken together, the results suggest that greater SG of gamma within the SMA (i.e., lower SMA Gamma GR thus exaggerated SG) correlated with better attention/executive function (i.e., lower TMT-B time), however this did not fully offset age-related deficits that emerge. Contralateral S1 theta SG did not significantly mediate the relationship between age and TMT-B performance (*p* > 0.05). Additionally, TMT-A was not found to be correlated with SG (*p* > 0.05).

**Figure 4.**
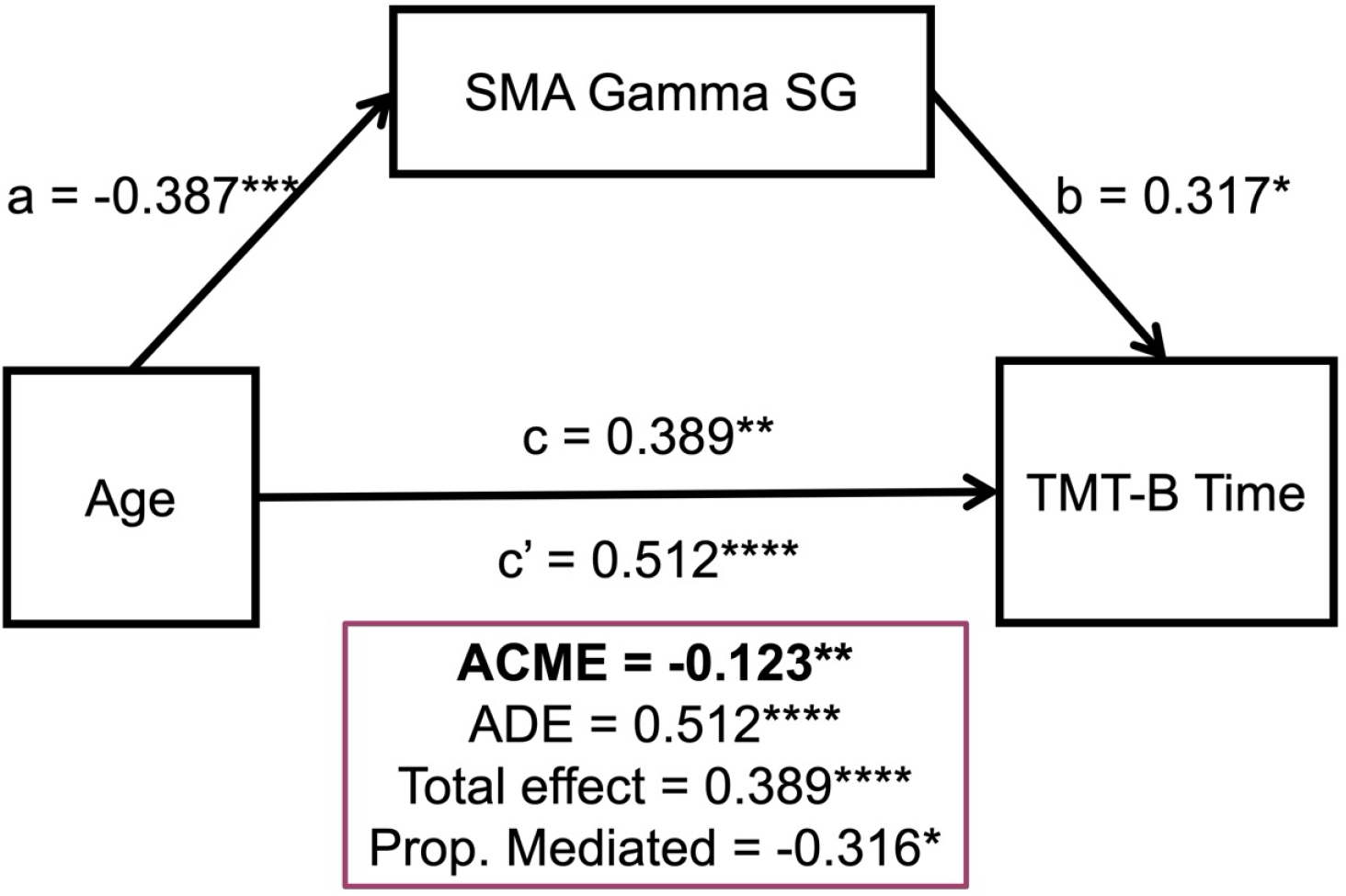
Mediation analysis of gamma (30-90 Hz) SG in contralateral SMA on attention/executive function and age. A partial, inconsistent mediation of TMT-B time on age through the mediator (i.e., SMA Gamma SG) was observed. Standardized regression coefficients are displayed. \**p* < 0.05; \*\**p* < 0.01; \*\*\**p* < 0.005; \*\*\*\**p* < 0.001. SG = somatosensory gating. SMA = supplementary motor area. TMT-B = trail making test part B. ACME = indirect effect. ADE = direct effect.

Lastly, structural and diffusion MRI analyses did not reveal any significant associations (see Supplementary Information for full results).

## Discussion

In the present study, we investigated the impact of healthy aging on somatosensory gating, a neural inhibitory mechanism, using a whole brain approach across numerous frequency bands. We first identified that theta SG was significantly associated with age in contralateral (left) S1; specifically, it was exaggerated with advancing age (i.e., more inhibition). We further demonstrated that the absolute amplitude of the S1 theta response following Stim2 significantly decreased with advancing age beyond the effects of baseline, lending to larger differences between Stim1 and Stim2 in older-aged adults. In the theta-band, we also identified decreases in spontaneous activity within contralateral S1 in older adults. Additionally, gamma SG was significantly impacted by age in the contralateral SMA, such that it was exaggerated with advancing age (i.e., greater inhibition). Further, we demonstrated that the relative amplitude of the SMA gamma response following Stim1 significantly increased with age, lending to larger differences between Stim1 and Stim2 in older-aged adults, compared to middle-aged adults. Finally, we report that the gating of gamma within SMA partially mediated age effects on TMT-B time, a measure of attention/executive function. These findings expand our current understanding of age-related SG processes and further highlight the complex interplay between sensory inhibition and higher-order cognition in an ethnoracially-diverse healthy aging population.

Novel to our study, we focused on multiple frequency bands of interest to SG (i.e., theta, alpha, beta, and gamma) and pursued a whole-brain approach. We found robust age-related gamma SG in regions outside of S1 (i.e., SMA) and age-related theta SG in S1. Interestingly, the gamma findings suggest age-related exaggeration of gamma SG in the higher-order sensory processing region, SMA. SMA has not been studied in the context of gamma SG. However, converging evidence indicates that SMA, a medial frontal region, supports motor control through gamma dynamics, particularly during sequential processing of motor and cognitive tasks [27]. For example, Chen et al. demonstrated that gamma activity in SMA was primarily modulated by proactive motor control (i.e., trial-dependent arm movement reaction time) [28]. In addition, Miyaguchi et al. showed that externally driving SMA at gamma frequency modulates bimanual motor performance, supporting a causal role for SMA gamma oscillations in motor coordination [57]. Together, these findings may suggest that SMA gamma reflects higher-order motor control processes rather than primary sensory encoding. Direct evidence on SMA gamma and aging remains limited. However, gamma oscillations are tightly linked to GABAergic inhibitory function [58–60], and age-related reductions in SMA inhibitory tone have been reported, with evidence suggesting that reduced SMA GABA may in fact support motor performance in older adults [61]. In this context, our finding of exaggerated gamma SG and increased gamma Stim1 responses in SMA of older adults may reflect age-related shifts in inhibitory regulation within higher-order motor networks.

Additionally, a plethora of past studies observed increased TMT-B time with age (i.e., declining attention/executive function) [23, 24], a pattern we also replicated. TMT-B is a time-sensitive measure indexing multiple cognitive domains, including attention, inhibition, executive function, visual perception, and motor speed, and is widely used to assess cognitive decline [33, 34]. Extending this literature, we found that SMA gamma SG inconsistently mediated the relationship between age and TMT-B performance, such that age effects on TMT-B increased after including SMA gamma SG in the model [56]. On the other hand, exaggerated SMA gamma SG (i.e., greater inhibition) was associated with better performance (lower completion time). Within the framework of cognitive aging, this pattern is consistent with partial compensatory recruitment, in which age-related changes in lower-level sensory processing are offset by increased engagement of higher-order regions, such as the SMA. In other words, exaggerated SMA gamma SG may reflect an adaptive shift toward greater top-down motor and sensory integration to support task performance. Importantly, however, the direct relationship between age and TMT-B performance remained significant even after accounting for SMA gamma SG, suggesting that this compensatory mechanism is beneficial but insufficient to fully counteract age-related declines in attention/executive function. SMA gamma SG accounted for 32% of variance, while 68% of the age-cognition relationship was not explained by it. With that said, the partial mediation is still highly informative, identifying SMA gamma SG as a functionally relevant neural correlate of preserved attention/executive function in aging, and suggesting that individuals who show greater age-related redistribution of inhibitory gating toward higher-order sensorimotor regions may derive a cognitive benefit from this reorganization. Further, the absence of an association between SG and TMT-A, alongside the observed relationship with TMT-B, suggests that SG may be more closely related to higher-order executive processes such as cognitive flexibility and inhibitory control, which are more specifically engaged during TMT-B performance. Together, these findings support a model in which age-related changes in SG are redistributed toward higher-order regions, with partial functional compensation for neurocognitive decline.

In contralateral S1, our findings indicate exaggerated theta SG with increasing age. To our knowledge, theta SG has rarely been examined. Key evidence comes from Wiesman et al., who observed theta SG in contralateral S1 and showed that it was enhanced when attention was directed toward somatosensory input relative to distraction, suggesting that early theta gating is sensitive to attentional state [4]. Given this, it is notable that theta SG did not mediate the relationship between age and attention/executive function herein, which may reflect the fact that Wiesman et al. tested within-subject attentional modulation of somatosensory inhibition, whereas our study asked whether age-related SG differences explain age-related differences in attention/executive function. More broadly, theta oscillations are consistently implicated in sensorimotor control and the coordination of large-scale networks [19, 20, 62–68], and multiple studies suggest that theta dynamics are age-sensitive [21, 69, 70]. Rempe et al. employed large-scale lifespan investigations into whole brain oscillatory dynamics and found age-related decreases in spontaneous theta power [69], consistent with our finding of reduced S1 theta baseline amplitude with age. Other studies investigating theta power in the brain have found region-specific decreases with age during memory tasks, and at rest, increased theta power is related to better cognitive performance on multiple domains (i.e., attention, verbal recall, executive function) [21, 70]. While the present study focused on investigating ties between SG and attention/executive function, probing whether theta SG and spontaneous theta have implications for age-related changes in other cognitive domains warrants consideration and may aid in interpretation of the functional relevance of our age effects seen herein. For example, our S1 theta Stim2 findings, where absolute Stim2 response amplitude declines beyond baseline effects while Stim1 does not, may reflect age-related changes in repetitive-stimulation responsiveness (e.g., increased inhibition and/or neural fatigue during the second stimulus), rather than a compensatory mechanism per se. Future studies that explicitly manipulate attention (as in Wiesman et al.) and incorporate neurochemical or microstructural measures will be important for distinguishing whether age-related theta SG reflects state-dependent attentional control versus age-related physiological changes in somatosensory circuitry.

It is important to note the limitations of our study. First, we did not replicate gamma SG deficits in S1 with age. Studies reporting this finding included participants in their 20s and early- and mid-30s [6, 18], however, we capture a middle-to older-aged healthy aging sample. As such, our restricted age range (39-78 years) may have limited sensitivity to detect earlier-life changes in S1 gamma SG. Additionally, the observed association between S1 gamma SG and age was characterized by a small effect size (η^2^ = 0.02), suggesting that subtle effects may have been underpowered in the current sample. Secondary and more broadly, the cross-sectional design precludes conclusions about within-subject aging trajectories, and future longitudinal work spanning a wider age range will be critical to fully characterize lifespan changes in somatosensory gating. Third, only TMT-A/B were used to index attention/executive function, and although TMT-B is tightly linked to inhibition, using a combination of neurocognitive tests is preferred. Additional neurocognitive tests for these domains can further inform the SG and attention/executive function relationship. Finally, TMT-B performance and SG relationship can also be influenced by motor performance; future studies with more statistical power can help to tease apart functional relationships between neuroimaging, neurocognitive, and behavioral (i.e., motor control or peripheral function) metrics.

The primary goal of the current study was to examine age-related effects on somatosensory gating across theta, alpha, beta, and gamma frequencies using a whole-brain voxel-wise approach. We found that gating of early sensory oscillatory dynamics (i.e., specifically theta and gamma) exhibits age-related increases in sensorimotor regions relevant to somatosensory perception and integration. Notably, SG was exaggerated with advancing age in lower-frequency bands (theta) within S1 and in higher-frequency bands (gamma) within higher-order regions such as the supplementary motor area (SMA). Given that theta oscillations are primarily known for network-wide and multisensory integration and learning, and gamma oscillations in sensory regions reflect basic perception [19–21, 66, 71–76], these findings suggest an age-related shift in inhibitory processing from primary sensory regions toward higher-order integrative regions. Importantly, exaggerated SMA gamma SG was associated with better cognitive performance, supporting a compensatory framework in which higher-order regions are increasingly recruited to maintain function in the context of age-related neural changes. However, this compensation appears partial, as it does not fully offset age-related declines in attention/executive function. These findings raise the possibility that age-related changes in SG reflect adaptations to underlying neurobiological alterations, such as shifts in inhibitory neurotransmission. Future studies incorporating complementary neuroimaging approaches, such as magnetic resonance spectroscopy or positron emission tomography, will be important for probing the neurochemical basis of SG, particularly with respect to GABAergic systems. Additionally, greater emphasis on theta SG in aging populations is warranted, as our findings highlight its potential functional relevance. Together, this work advances our understanding of how somatosensory inhibition is reorganized with age and provides a foundation for future investigations into the neural mechanisms supporting healthy cognitive aging.

## Acknowledgements

We would like to acknowledge Adriana Ohm in her assistance with DHMS piloting and data collection.

## Data availability

Requests for access to DHMS data may be submitted to the Dallas Hearts Study (DHS) Publications Committee according to established study procedures. The code used for analyses in this study are available from the corresponding author upon reasonable request.

## Author Contributions

ALP and EMD contributed to the development, design, and validation of the MEG methodology. FFY contributed to the development and design of the MRI methodology. LHL, HR, and CMC contributed to the development and design of the neuropsychological methodology. MV completed the formal MEG data processing, analyses and investigation, and wrote the original draft of the manuscript. MV, LG, and SW contributed to MRI data preprocessing, and ALP, EMD, MV, NMB, LG, and SW assisted in data collection. YX, FFY, AMS, and ALP contributed to the supervision of the data analyses. YX, NMB, TP, LG, SW, FFY, LHL, HR, CMC, AMS, EMD, JAM, and ALP provided edits to the manuscript.

## References

1. Commodari, E. and M. Guarnera, Attention and aging. Aging Clin Exp Res, 2008. 20(6): p. 578–84.

2. Harada, C.N., M.C. Natelson Love, and K.L. Triebel, Normal cognitive aging. Clin Geriatr Med, 2013. 29(4): p. 737–52.

3. Cromwell, H.C., et al., Sensory gating: a translational effort from basic to clinical science. Clin EEG Neurosci, 2008. 39(2): p. 69–72.

4. Wiesman, A.I. and T.W. Wilson, Attention modulates the gating of primary somatosensory oscillations. Neuroimage, 2020. 211: p. 116610.

5. Wiesman, A.I., et al., Oscillatory dynamics and functional connectivity during gating of primary somatosensory responses. J Physiol, 2017. 595(4): p. 1365–1375.

6. Spooner, R.K., et al., Rhythmic Spontaneous Activity Mediates the Age-Related Decline in Somatosensory Function. Cereb Cortex, 2019. 29(2): p. 680–688.

7. Spooner, R.K., et al., Methodological considerations for a better somatosensory gating paradigm: The impact of the inter-stimulus interval. Neuroimage, 2020. 220: p. 117048.

8. Arif, Y., et al., Aberrant inhibitory processing in the somatosensory cortices of cannabis-users. J Psychopharmacol, 2021. 35(11): p. 1356–1364.

9. Spooner, R.K., et al., Aberrant oscillatory dynamics during somatosensory processing in HIV-infected adults. Neuroimage Clin, 2018. 20: p. 85–91.

10. Liu, Y.T., et al., Aberrant Sensory Gating of the Primary Somatosensory Cortex Contributes to the Motor Circuit Dysfunction in Paroxysmal Kinesigenic Dyskinesia. Front Neurol, 2018. 9: p. 831.

11. Trevarrow, M.P., et al., Altered Somatosensory Cortical Activity Is Associated with Cortical Thickness in Adults with Cerebral Palsy: Multimodal Evidence from MEG/sMRI. Cereb Cortex, 2022. 32(6): p. 1286–1294.

12. Kurz, M.J., et al., Children with Cerebral Palsy Hyper-Gate Somatosensory Stimulations of the Foot. Cereb Cortex, 2018. 28(7): p. 2431–2438.

13. Casagrande, C.C., et al., Impact of HIV-infection on human somatosensory processing, spontaneous cortical activity, and cortical thickness: A multimodal neuroimaging approach. Hum Brain Mapp, 2021. 42(9): p. 2851–2861.

14. Spooner, R.K., et al., Prefrontal gating of sensory input differentiates cognitively impaired and unimpaired aging adults with HIV. Brain Commun, 2020. 2(2): p. fcaa080.

15. Arpin, D.J., et al., A reduced somatosensory gating response in individuals with multiple sclerosis is related to walking impairment. J Neurophysiol, 2017. 118(4): p. 2052–2058.

16. Casagrande, C.C., et al., Signatures of somatosensory cortical dysfunction in Alzheimer’s disease and HIV-associated neurocognitive disorder. Brain Commun, 2022. 4(4): p. fcac169.

17. Wiesman, A.I., et al., Somatosensory dysfunction is masked by variable cognitive deficits across patients on the Alzheimer’s disease spectrum. EBioMedicine, 2021. 73: p. 103638.

18. Proskovec, A.L., et al., Local cortical thickness predicts somatosensory gamma oscillations and sensory gating: A multimodal approach. Neuroimage, 2020. 214: p. 116749.

19. Fries, P., Rhythms for Cognition: Communication through Coherence. Neuron, 2015. 88(1): p. 220–35.

20. Fan, J., et al., The relation of brain oscillations to attentional networks. J Neurosci, 2007. 27(23): p. 6197–206.

21. Finnigan, S. and I.H. Robertson, Resting EEG theta power correlates with cognitive performance in healthy older adults. Psychophysiology, 2011. 48(8): p. 1083–7.

22. Dockstader, C., D. Cheyne, and R. Tannock, Cortical dynamics of selective attention to somatosensory events. Neuroimage, 2010. 49(2): p. 1777–85.

23. Rasmusson, X.D., Zonderman, A. B., Kawas, C., & Resnick, S. M., Effects of Age and Dementia on the Trail Making Test. The Clinical Neuropsychologist, 1998. 12(2): p. 169–178.

24. Kennedy, K.J., Age effects on Trail Making Test performance. Percept Mot Skills, 1981. 52(2): p. 671–5.

25. Arif, Y., et al., Modulation of attention networks serving reorientation in healthy aging. Aging (Albany NY), 2020. 12(13): p. 12582–12597.

26. Mima, T., et al., Somesthetic function of supplementary motor area during voluntary movements. Neuroreport, 1999. 10(9): p. 1859–62.

27. Cona, G. and C. Semenza, Supplementary motor area as key structure for domain-general sequence processing: A unified account. Neurosci Biobehav Rev, 2017. 72: p. 28–42.

28. Chen, X., K.W. Scangos, and V. Stuphorn, Supplementary motor area exerts proactive and reactive control of arm movements. J Neurosci, 2010. 30(44): p. 14657–75.

29. Bell, N.M., et al., Magnetoencephalography in human neuroscience research: planning, piloting, implementation and quality assurance. Nat Protoc, 2026.

30. Hingson, R.W., W. Zha, and A.M. White, Drinking Beyond the Binge Threshold: Predictors, Consequences, and Changes in the U.S. Am J Prev Med, 2017. 52(6): p. 717–727.

31. Avila-Villanueva, M. and M.A. Fernandez-Blazquez, Subjective Cognitive Decline as a Preclinical Marker for Alzheimer’s Disease: The Challenge of Stability Over Time. Front Aging Neurosci, 2017. 9: p. 377.

32. Rabin, L.A., C.M. Smart, and R.E. Amariglio, Subjective Cognitive Decline in Preclinical Alzheimer’s Disease. Annu Rev Clin Psychol, 2017. 13: p. 369–396.

33. Kortte, K.B., M.D. Horner, and W.K. Windham, The trail making test, part B: cognitive flexibility or ability to maintain set? Appl Neuropsychol, 2002. 9(2): p. 106–9.

34. Llinas-Regla, J., et al., The Trail Making Test. Assessment, 2017. 24(2): p. 183–196.

35. Reitan, R.M., Validity of the Trail Making Test as an indicator of organic brain damage. Perceptual and Motor Skills, 1958. 8: p. 271–276.

36. Stephen, J.M., et al., Somatosensory responses in normal aging, mild cognitive impairment, and Alzheimer’s disease. J Neural Transm (Vienna), 2010. 117(2): p. 217–25.

37. Zhang, J., et al., Gender differences of neuropsychological profiles in cognitively normal older people without amyloid pathology. Compr Psychiatry, 2017. 75: p. 22–26.

38. Gross, A.L., et al., Effects of education and race on cognitive decline: An integrative study of generalizability versus study-specific results. Psychol Aging, 2015. 30(4): p. 863–880.

39. Posit, RStudio: Integrated Development Environment for R. 2025, Posit Software, PBC: Boston, MA.

40. Medvedovsky, M., et al., Artifact and head movement compensation in MEG. Neurol Neurophysiol Neurosci, 2007: p. 4.

41. Taulu, S. and J. Simola, Spatiotemporal signal space separation method for rejecting nearby interference in MEG measurements. Phys Med Biol, 2006. 51(7): p. 1759–68.

42. Taulu, S., M. Kajola, and J. Simola, Suppression of interference and artifacts by the Signal Space Separation Method. Brain Topogr, 2004. 16(4): p. 269–75.

43. Tadel, F., et al., Brainstorm: a user-friendly application for MEG/EEG analysis. Comput Intell Neurosci, 2011. 2011: p. 879716.

44. Fonov, V.S., et al., Unbiased nonlinear average age-appropriate brain templates from birth to adulthood. Neuroimage, 2009. 47: p. S102.

45. Van Veen, B.D., et al., Localization of brain electrical activity via linearly constrained minimum variance spatial filtering. IEEE Trans Biomed Eng, 1997. 44(9): p. 867–80.

46. Hillebrand, A., et al., A new approach to neuroimaging with magnetoencephalography. Hum Brain Mapp, 2005. 25(2): p. 199–211.

47. Liljestrom, M., et al., Neuromagnetic localization of rhythmic activity in the human brain: a comparison of three methods. Neuroimage, 2005. 25(3): p. 734–45.

48. Statistical Parametric Mapping: The Analysis of Functional Brain Images. 2006. 688.

49. Worsley, K.J., et al., Detecting changes in nonisotropic images. Hum Brain Mapp, 1999. 8(2-3): p. 98–101.

50. Poline, J.B., et al., Estimating smoothness in statistical parametric maps: variability of p values. J Comput Assist Tomogr, 1995. 19(5): p. 788–96.

51. Worsley, K.J., et al., A unified statistical approach for determining significant signals in images of cerebral activation. Hum Brain Mapp, 1996. 4(1): p. 58–73.

52. MacKinnon, D.P., A.J. Fairchild, and M.S. Fritz, Mediation analysis. Annu Rev Psychol, 2007. 58: p. 593–614.

53. Baron, R.M. and D.A. Kenny, The moderator-mediator variable distinction in social psychological research: conceptual, strategic, and statistical considerations. J Pers Soc Psychol, 1986. 51(6): p. 1173–82.

54. Dustin Tingley, T.Y., Kentaro Hirose, Luke Keele, Kosuke Imai, mediation: R Package for Causal Mediation Analysis. Journal of Statistical Software, 2014. 59(5): p. 1–38.

55. Rossetti, H.C., et al., Normative data for the Montreal Cognitive Assessment (MoCA) in a population-based sample. Neurology, 2011. 77(13): p. 1272–5.

56. MacKinnon, D.P., J.L. Krull, and C.M. Lockwood, Equivalence of the mediation, confounding and suppression effect. Prev Sci, 2000. 1(4): p. 173–81.

57. Miyaguchi, S., et al., Effects of stimulating the supplementary motor area with a transcranial alternating current for bimanual movement performance. Behav Brain Res, 2020. 393: p. 112801.

58. Gaetz, W., et al., Relating MEG measured motor cortical oscillations to resting gamma-aminobutyric acid (GABA) concentration. Neuroimage, 2011. 55(2): p. 616–21.

59. Kujala, J., et al., Gamma oscillations in V1 are correlated with GABA(A) receptor density: A multi-modal MEG and Flumazenil-PET study. Sci Rep, 2015. 5: p. 16347.

60. Muthukumaraswamy, S.D., et al., Resting GABA concentration predicts peak gamma frequency and fMRI amplitude in response to visual stimulation in humans. Proc Natl Acad Sci U S A, 2009. 106(20): p. 8356–61.

61. Frieske, J., et al., Age-related GABA- and glutamatergic differences in SMA during bimanual coordination. Imaging Neurosci (Camb), 2025. 3.

62. Begus, K. and E. Bonawitz, The rhythm of learning: Theta oscillations as an index of active learning in infancy. Dev Cogn Neurosci, 2020. 45: p. 100810.

63. Darna, M., et al., Frontal theta oscillations and cognitive flexibility: Age-related modulations in EEG activity. Aging Brain, 2025. 8: p. 100142.

64. Michels, L., et al., Simultaneous EEG-fMRI during a working memory task: modulations in low and high frequency bands. PLoS One, 2010. 5(4): p. e10298.

65. Nyhus, E. and T. Curran, Functional role of gamma and theta oscillations in episodic memory. Neurosci Biobehav Rev, 2010. 34(7): p. 1023–35.

66. Summerfield, C. and J.A. Mangels, Coherent theta-band EEG activity predicts item-context binding during encoding. Neuroimage, 2005. 24(3): p. 692–703.

67. Vlahou, E.L., et al., Resting-state slow wave power, healthy aging and cognitive performance. Sci Rep, 2014. 4: p. 5101.

68. Wang, X.J., Neurophysiological and computational principles of cortical rhythms in cognition. Physiol Rev, 2010. 90(3): p. 1195–268.

69. Rempe, M.P., et al., Spontaneous cortical dynamics from the first years to the golden years. Proc Natl Acad Sci U S A, 2023. 120(4): p. e2212776120.

70. Cummins, T.D. and S. Finnigan, Theta power is reduced in healthy cognitive aging. Int J Psychophysiol, 2007. 66(1): p. 10–7.

71. Fries, P., D. Nikolic, and W. Singer, The gamma cycle. Trends Neurosci, 2007. 30(7): p. 309–16.

72. Wulff-Abramsson, A., et al., Event-related theta synchronization over sensorimotor areas differs between younger and older adults and is related to bimanual motor control. Neuroimage, 2025. 308: p. 121032.

73. Gu, Z., et al., Hippocampus and Entorhinal Cortex Recruit Cholinergic and NMDA Receptors Separately to Generate Hippocampal Theta Oscillations. Cell Rep, 2017. 21(12): p. 3585–3595.

74. Hauck, M., J. Lorenz, and A.K. Engel, Attention to painful stimulation enhances gamma-band activity and synchronization in human sensorimotor cortex. J Neurosci, 2007. 27(35): p. 9270–7.

75. Gross, J., et al., Gamma oscillations in human primary somatosensory cortex reflect pain perception. PLoS Biol, 2007. 5(5): p. e133.

76. Rossiter, H.E., et al., Gamma oscillatory amplitude encodes stimulus intensity in primary somatosensory cortex. Front Hum Neurosci, 2013. 7: p. 362.

