## Supplemental Information for "Age-Related Alterations in Multispectral Somatosensory Gating: Evidence for Partial Compensation in Attentional Performance"

Supplementary Information

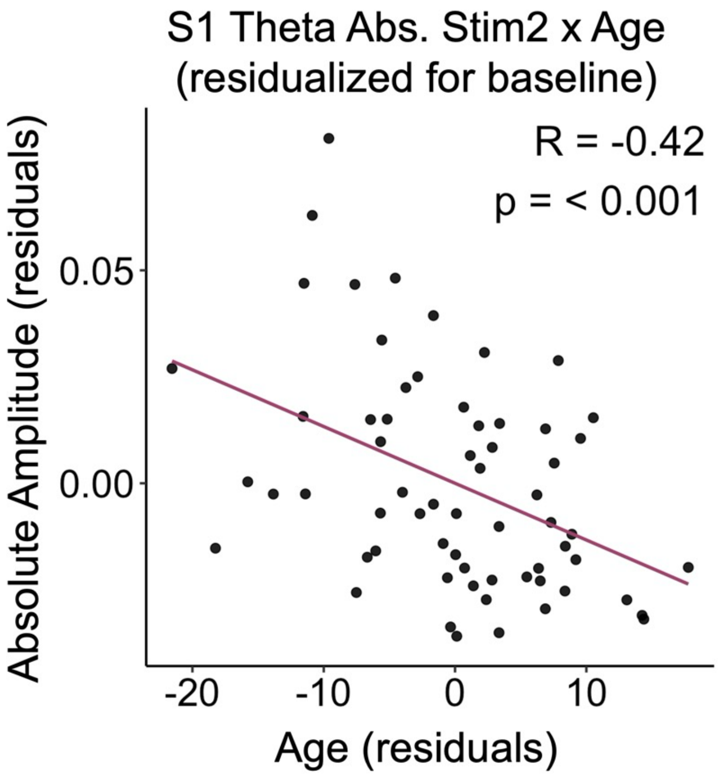


Supplemental Figure 1. Age Effects on Absolute Theta Stim2 Response Amplitude Independent of Theta Baseline. Age was negatively correlated with contralateral S1 absolute theta Stim2 response amplitude, above and beyond the effects of spontaneous theta amplitude in the region. S1 = primary somatosensory cortex.

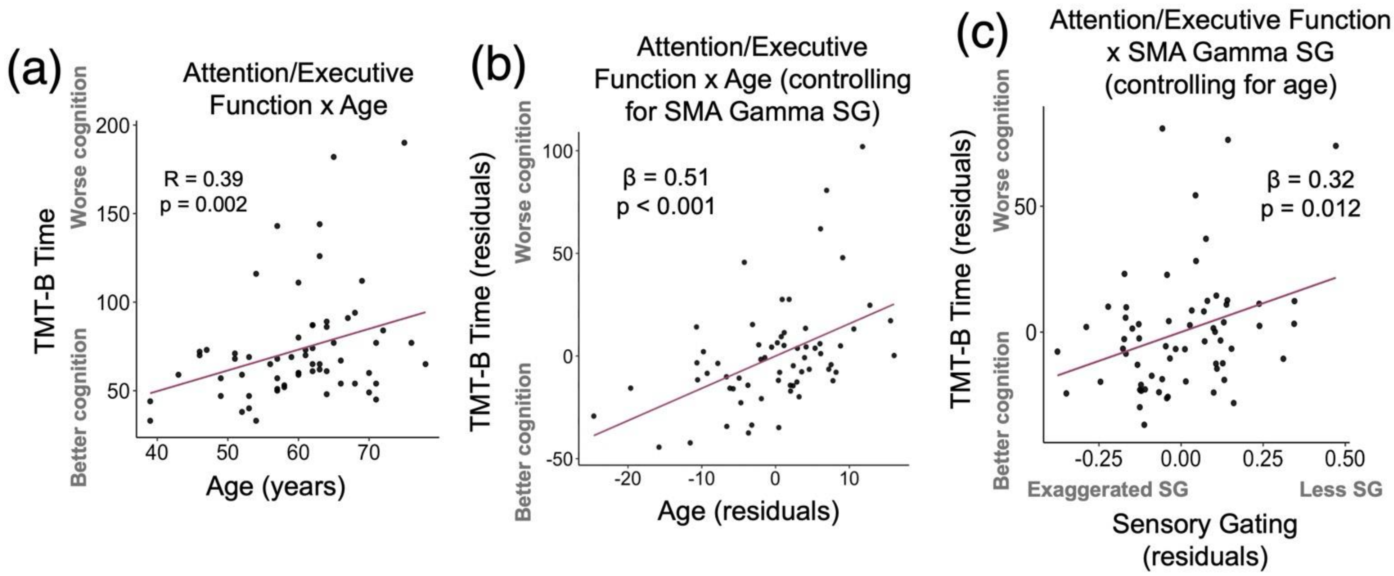


Supplemental Figure 2. Trail B, Age, and SMA Gamma SG Regression Plots. (a) Age was positively associated with Trail Making Test Part B (TMT-B) time, such that, as age increased, demographically adjusted TMT-B time increased (i.e., worse cognition/slower performance). TMT-B time is depicted on the y-axis, with higher values indicating worse cognition. Age in years is depicted on the x-axis. (b) A residual plot depicting attention/executive function and age relationship after controlling for SMA gamma SG. The y-axis depicts TMT-B time after removing SMA gamma SG variance. The x-ais displays residualized age, after controlling for SMA gamma SG. (c) A residual plot depicting TMT-B time and contralateral SMA gamma SG relationship after controlling for the effects of age. SMA gamma SG was positively associated with TMT-B time, such that, with greater gamma SG exaggerated, TMT-B time decreased (i.e., better cognition/faster performance). The y-axis represents TMT-B time after accounting for age. The x-axis represents SMA gamma SG after accounting for age.

Supplementary methodology for surface-based morphometry analyses

Since past studies have highlighted relationships between cortical thickness in the postcentral gyrus (corresponding to S1) and SG [1], we investigated whether S1 surface-based morphometry was related to age and MEG metrics.

Cortical thickness and gray matter (GM) volume were quantified using the automated recon-all processing stream in FreeSurfer (version 7.1.1) [2]. This pipeline includes steps such as intensity normalization, skull stripping, segmentation of white and gray matter, and reconstruction of cortical surfaces from each participant's high-resolution T1-weighted MRI. Cortical thickness was outlined as the distance between the white matter boundary and the pial surface at each vertex, whereas GM volume was derived from the parcellated anatomical outputs generated during processing. Vertex-wise estimates of thickness and volume were mapped onto anatomical regions using the Desikan-Killiany-Tourville atlas, from which region-of-interest (ROI) values were extracted for the left (contralateral) postcentral gyrus. Intracranial volume was included as a covariate in all analyses involving volumetric measures. ROI-derived cortical thickness and GM volume values were entered into linear regression models to examine associations with age. In additional exploratory analyses, these structural measures were also assessed in relation to MEG-derived somatosensory indices; however, no significant relationships were observed (all *p*'s $< 0.05$).

Supplementary methodology for diffusion analyses using fractional anisotropy

Past studies have suggested that fractional anisotropy (FA) changes as a function of age and cognitive health, specifically white matter tracts adjacent to sensorimotor regions were negatively impacted with age and correlated with poorer cognitive performance in attention/executive function domains [3]. Thus, we evaluated whether FA changes as a function of healthy cognitive aging and probed relationships with SG using tract-based spatial statistics (TBSS).

Diffusion-weighted imaging data were collected using a 102-direction multi-shell acquisition with b-values of 0, 500, 1000, 2000, and 3000 s/mm², along with an additional reversed phase-encoding b0 volume to enable correction of susceptibility-related distortions. Preprocessing was conducted using the FMRIB Software Library (FSL; version 6.1), including distortion correction with topup and correction for eddy currents and head motion [4]. Diffusion tensor imaging (DTI) was performed; tensors were fitted at each voxel using FSL's dtifit, producing FA maps for each participant. TBSS was used to perform voxel-wise analysis of white matter integrity [5]. FA maps were nonlinearly aligned to the FMRIB58_FA template, skeletonized (FA > 0.2), and each participant's data were projected onto the resulting white matter skeleton [4, 5]. Statistical analyses were conducted using permutation-based nonparametric testing (5,000 permutations). Age was included as the primary predictor. Threshold-free cluster enhancement was applied, and multiple comparisons corrections utilized family-wise error at $p < 0.05$. To explore structure-function relationships, mean FA values from significant clusters were extracted and examined in relation to MEG-derived somatosensory measures. Additionally, secondary analyses included SG metrics as primary predictors and age as a nuisance variable. Overall, these MEG and DTI exploratory analyses did not reveal any significant associations (all voxel-wise $p$'s > 0.05).

References


1. Proskovec, A.L., et al., *Local cortical thickness predicts somatosensory gamma oscillations and sensory gating: A multimodal approach.* Neuroimage, 2020. **214**: p. 116749.
2. Fischl, B., *FreeSurfer.* Neuroimage, 2012. **62**(2): p. 774-81.
3. Grieve, S.M., et al., *Cognitive aging, executive function, and fractional anisotropy: a diffusion tensor MR imaging study.* AJNR Am J Neuroradiol, 2007. **28**(2): p. 226-35.
4. Jenkinson, M., et al., *Fsl.* Neuroimage, 2012. **62**(2): p. 782-90.
5. Smith, S.M., et al., *Tract-based spatial statistics: voxelwise analysis of multi-subject diffusion data.* Neuroimage, 2006. **31**(4): p. 1487-505.
